# SARS-CoV-2 infection of primary human lung epithelium for COVID-19 modeling and drug discovery

**DOI:** 10.1101/2020.06.29.174623

**Authors:** A. Mulay, B. Konda, G. Garcia, C. Yao, S. Beil, C. Sen, A. Purkayastha, J. K. Kolls, D. A. Pociask, P. Pessina, J. Sainz de Aja, C. Garcia-de-Alba, C. F. Kim, B. Gomperts, V. Arumugaswami, B.R. Stripp

## Abstract

Coronavirus disease 2019 (COVID-19) is the latest respiratory pandemic resulting from zoonotic transmission of severe acute respiratory syndrome-related coronavirus 2 (SARS-CoV-2). Severe symptoms include viral pneumonia secondary to infection and inflammation of the lower respiratory tract, in some cases causing death. We developed primary human lung epithelial infection models to understand responses of proximal and distal lung epithelium to SARS-CoV-2 infection. Differentiated air-liquid interface cultures of proximal airway epithelium and 3D organoid cultures of alveolar epithelium were readily infected by SARS-CoV-2 leading to an epithelial cell-autonomous proinflammatory response. We validated the efficacy of selected candidate COVID-19 drugs confirming that Remdesivir strongly suppressed viral infection/replication. We provide a relevant platform for studying COVID-19 pathobiology and for rapid drug screening against SARS-CoV-2 and future emergent respiratory pathogens.

**One Sentence Summary:** A novel infection model of the adult human lung epithelium serves as a platform for COVID-19 studies and drug discovery.

## Introduction

COVID-19 caused by infection with the SARS-CoV-2 virus, is the most recent of a series of severe viral infections that typically initiate in the upper respiratory tract but have the potential to cause life-threatening pneumonia due to infection and inflammation of the lower respiratory tract (*1, 2*). Unlike human coronaviruses that lead to a self-limiting upper respiratory tract infection, SARS-CoV-2 and related viruses SARS-CoV and MERS-CoV are thought to originate from bats and cause severe symptoms in their human hosts due to lack of host-pathogen adaptation (*3*). Similar zoonotic transmission of other respiratory pathogens, such as H1N1 influenza A virus (*4*) are associated with recurrent pandemics with severe pulmonary complications seen among infected individuals due to infection of the lower respiratory tract. Even though acute viral pneumonia appears to be a common pathological outcome of infection with these severe respiratory viruses, mechanisms leading to these adverse outcomes are poorly understood. We sought to develop culture models of proximal and distal human lung epithelium as a platform to define disease mechanisms and for rapid drug discovery in the setting of current and future severe respiratory viral infections. Moreover, this study is the first report of a primary *in vitro* system of the adult human alveoli to model COVID-19.

## Materials and Methods

Human lung tissue was obtained from deceased organ donors in compliance with consent procedures developed by International Institute for the Advancement of Medicine (IIAM) and approved by the Internal Review Board at Cedar-Sinai Medical Center.

### Cell isolation

Human lung tissue was processed as described previously (*5–7*) with the following modifications. For isolation of proximal airway cells, trachea and the first 2-3 generation of bronchi were slit vertically and enzymatically digested with Liberase (50 μg/mL) and DNase 1 (25 μg/mL) incubated at 37°C with mechanical agitation for 20 minutes, followed by gentle scraping of epithelial cells from the basement membrane. The remaining tissue was finely minced and further digested for 40 minutes at 37°C. For distal alveolar cell isolation, small airways of 2mm diameter or less and surrounding parenchymal tissue was minced finely and enzymatically digested for 40-60 minutes as described before. Total proximal or distal dissociated cells were passed through a series of cell strainers of decreasing pore sizes from 500μm to 40μm under vacuum pressure and depleted of immune and endothelial cells by magnetic associated cell sorting (MACS) in accordance to the manufacturing protocol (Miltenyi Biotec). Viable epithelial cells were further enriched by fluorescence associated cell sorting (FACS) using DAPI (Thermo Fisher Scientific) and antibodies against EPCAM (CD326), CD45 and CD31 (Biolegend) on a BD Influx cell sorter (Becton Dickinson).

### Culture and differentiation of proximal airway epithelial cells at air liquid interface

FACS enriched proximal airway epithelial cells were expanded in T25 or T75 flasks coated with bovine type I collagen (Purecol, Advanced biomatrix) in Pneumacult Ex media (STEMCELL Technologies), supplemented with 1X Penicillin-Streptomycin-Neomycin (PSN) Antibiotic Mixture (Thermo Fisher Scientific) and 10μM Rho kinase inhibitor, Y-27632 (STEMCELL technologies). Upon confluence cells were dissociated using 0.05% Trypsin-EDTA (Thermo Fisher Scientific) and seeded onto collagen coated 0.4 μm pore size transparent cell culture inserts in a 24-well supported format (Corning) at a density of 7.5 × 10^4^ cells per insert. Cells were initially cultured submerged with 300 μL of Pneumacult Ex media in the apical chamber and 700 μL in the basement chamber for 3-5 days. Upon confluence, cells were cultured at air liquid interface in 700 μL Pneumacult ALI media (STEMCELL Technologies) supplemented with 1X PSN and media was changed every 48 hrs. Cultures were maintained at 37°C in a humidified incubator (5% CO_2_) and used for SARS-CoV-2 infection after 16-20 days of differentiation.

### Culture of 3D alveolar organoids

Five thousand FACS enriched distal lung epithelial cells were mixed with 7.5 × 10^4^ MRC5 human lung fibroblast cells (ATCC CCL-171) and resuspended in a 50:50 (v/v) ratio of ice cold Matrigel (Corning) and Pneumacult ALI medium. 100uL of the suspension was seeded onto the apical surface of a 0.4 μm pore-size cell culture insert in a 24 well supported format. After polymerization of Matrigel, 700μL of Pneumacult ALI medium was added to the basement membrane. Media was supplemented with 50μg per ml of Gentamycin (Sigma Aldrich) for the first 24 hrs. and 10μM Rho kinase inhibitor for the first 48 hrs. 2 μM of the Wnt pathway activator, CHIR-99021 (STEMCELL technologies) was added to the media at 48hrs and maintained for the entire duration of culture. Media was changed every 48 hrs. Cultures were maintained at 37°C in a humidified incubator (5% CO_2_) and used for SARS-Co-V2 infection after 15 days.

### SARS-CoV-2 infection of ALI and organoid cultures

All studies involving SARS-CoV-2 infection of proximal and distal lung epithelial cells were conducted in the UCLA BSL3 High-Containment Facility. SARS-CoV-2, isolate USA-WA1/2020, was sourced from the Biodefense and Emerging Infections (BEI) Resources of the National Institute of Allergy and Infectious Diseases (NIAID). SARS-CoV-2 was amplified once in Vero-E6 cells and viral stocks were stored at −80°C. Vero-E6 cells were cultured in Eagle’s minimal essential medium (MEM) (Corning) supplemented with 10% fetal bovine serum (Corning), penicillin-streptomycin (100 units/ml, Gibco), 2 mM L-glutamine (Gibco) and 10 mM HEPES (Gibco). Cells were incubated at 37°C in a humidified incubator (5% CO_2_).

Prior to infection of alveolar organoid cultures, Matrigel was dissolved by adding 500μL of Dispase (500μg/ml, Gibco) to the apical and basement chambers of inserts and incubating for 1 hour at 37°C. Organoids were harvested, washed, gently triturated with a P1000 tip by pipetting up and down. SARS-CoV-2 inoculum (1×10^4 TCID50 per well) was added to the alveolar organoids in a 2 ml conical tube and incubated for 2 hours at 37°C (5% CO2). Every 15 minutes, tubes were gently mixed to facilitate virus adsorption on to the cells. Subsequently, inoculum was replaced with fresh Pneumacult ALI medium and organoids were transferred to the apical chamber of the insert in 100 μL volume with 500 μL of the medium in the basement chamber. Cultures were incubated at 37°C (5% CO2) and harvested at indicated time points for sample collection.

For infection of proximal airway ALI cultures, 100 μl of SARS-CoV-2 virus inoculum (1×10^4 TCID50 per well) was added on to the apical chamber of inserts. Cells were incubated at 37°C (5% CO2) for 2 hours for virus adsorption. Subsequently, the cells were washed and fresh Pneumacult media (500 μl in the base chamber) was added. Cells were incubated at 37°C (5% CO2 and harvested for analysis at indicated time points for sample collection.

### Drug validation experiments

Hydroxychloroquine (Selleck Chemicals Cat. No. S4430) and Remdesivir (Selleck Chemicals Cat. No. S8932) were dissolved in DMSO to a stock concentration of 10 mM. IFNB1 stock of 10^6 units/ml was provided by Dr. Jay Kolls. To test the efficacy of the drugs against SARS-CoV-2 infection/replication in proximal airway epithelial cells and AT2 cells, cultures were treated with 10 μM of Hydroxychloroquine or Remdesivir, or 100 units/ml of IFNB1.

For treatment of proximal airway epithelial cells, Pneumacult ALI media containing drugs was added to ALI cultures (100 μL to the apical chamber and 700 μL to the basement chamber), 3 hours prior to infection. 100 μL of SARS-CoV-2 viral inoculum (1×10^4 TCID50 per well) was added to the apical chamber. After 2 hours of viral adsorption, cells were washed and 700 μL of fresh Pneumacult ALI media containing drugs was added to the basement chamber. Drugs were maintained in the media for the duration of culture post infection. ALI cultures without drug treatment in the presence or absence (mock) of viral infections were included as controls.

For alveolar organoid drug study, dissociated organoids were suspended in 100μL of Pneumacult ALI media containing drugs 3 hours prior to SARS-CoV-2 infection. Viral infections were performed as described previously. Drugs were maintained in the media for the duration of culture post infection. Organoid cultures without drug treatment in the presence or absence (mock) of viral infections were included as controls.

### Immunofluorescence staining

1-3 days after SARS-CoV-2 infection, cultures were fixed by adding 500μL of 4% paraformaldehyde in phosphate-buffered saline (PBS) for 20 minutes. Fixed samples were permeabilized and blocked for 1 hour in a ‘blocking buffer’ containing PBS, 2% bovine serum albumin, 5% goat serum, 5% donkey serum and 0.3% Triton X-100. Primary antibodies diluted in the blocking buffer were applied to samples and incubated overnight at 4°C. The following primary antibodies were used: Acetylated tubulin (1:200, Sigma Aldrich, Cat. No. T6793); Mucin 5AC, 45M1 (1:500, Thermo Fisher Scientific, Cat. No. MA5-12178); HTII-280 (1:500, Terrace Biotech, Cat. No. TB-27AHT2-280); SARS-CoV-2 spike (S) protein (1:100, BEI Resources NR-616 Monoclonal Anti-SARS-CoV S Protein (Similar to 240C) SARS coronavirus); Cleaved caspase-3 (1:200, Cell Signaling Technology Cat. No. 9661). Samples were washed 3 times for 5 minutes each with PBS. Appropriate secondary antibodies conjugated to fluorophores (Thermo Fisher Scientific) were applied to the samples for 1 hour at room temperature. Samples were washed 3 times for 5 minutes each with PBS followed by addition of DAPI (1:5000) for 5 minutes. Insert membranes were carefully detached from their transwell support with a fine scalpel, and transferred to a glass microscopic slide. Samples were mounted using Fluoromount G (Thermo Fisher Scientific), imaged on a Zeiss LSM 780 Confocal Microscope and images were processed using Zen Blue software (Zeiss).

### Real time quantitative PCR (RT-qPCR)

Total RNA was extracted from mock and SARS-CoV-2 infected proximal airway ALI and alveolar organoid cultures lysed in Trizol (Thermo Fisher Scientific) using the chloroform-iso-propanol-ethanol method. 500ng of RNA was reversed transcribed into cDNA in a 20 μL reaction volume using iScript cDNA synthesis kit (Biorad) in accordance to manufacturer’s guidelines. RT-qPCR was performed on 10ng of CDNA per reaction in triplicates for each sample using SYBR green master mix (Thermo Fisher Scientific) on a 7500 Fast Real Time PCR system (Applied biosystems). Primers sequences for detection of SARS-CoV-2 N gene were obtained from the Center for Disease Control’s resources for research labs. Primer pairs used are as follows:

2019-nCoV_N1 Forward: GAC CCC AAA ATC AGC GAA AT
2019-nCoV_N1 Reverse: TCT GGT TAC TGC CAG TTG AAT CTG
GAPDH Forward: CCACCTTTGACGCTGGG
GAPDH Reverse: CATACCAGGAAATGAGCTTGACA

### Bulk RNA sequencing

2 days after SARS-CoV-2 infection, total RNA was extracted for bulk RNA sequencing as described before. RNA quality was analyzed using the 2100 Bioanalyzer (Agilent Technologies) and quantified using QubitTM (ThermoFisher Scientific). Library construction was performed by the Cedars-Sinai Applied Genomics, Computation and Translational Core using the Lexogen RiboCop rRNA Depletion kit and Swift Biosciences RNA Library Sciences. Sequencing was performed using the the NovaSeq 6000 (Illumina) with single-end 75bp sequencing chemistry. On average, about 20 million reads were generated from each sample. Raw reads were aligned using Star aligner 2.6.1 (*8*)/RSEM 1.2.28 (*9*) with default parameters, using a custom human GRCh38 transcriptome reference downloaded from http://www.gencodegenes.org, containing all protein coding and long non-coding RNA genes based on human GENCODE version 33 annotation with SARS-Cov2 virus genome MT246667.1 https://www.ncbi.nlm.nih.gov/nuccore/MT246667.1.

Differential gene expression was determined by DESeq2 (*10*). Top differential genes were determined based on fold change and test statistics. Pathway analysis was performed using Ingenuity Pathway Analysis.

### Statistics

Statistical analysis was performed using GraphPad Prism version 8 software. Data is presented as linear fold change or log_2_ fold change ± SEM. SARS-CoV-2 infection time course for ALI and organoid cultures was analyzed using Two- Way ANOVA with Sidak’s post hoc correction. Drug validation experiments were analyzed using One-way ANOVA with Tukey’s post hoc correction.

## Results

### SARS-CoV-2 infects and replicates within human proximal airway epithelial cells

Since the upper respiratory tract represents the most likely initial site of respiratory virus infection, we initially utilized the well-established air-liquid interface (ALI) culture system to study the effect of SARS-CoV-2 infection on proximal airway epithelial cells (Figure 1A). Human trachea-bronchial epithelial cells (HBECs) isolated from trachea and upper bronchi were differentiated at ALI for 16-20 days to yield a pseudostratified mucociliary epithelium and infected with SARS-CoV-2 (1×10^4 TCID50 per well). Cells were harvested for analysis at 1 to 3 days post infection (dpi) (Figure 1A). SARS-CoV-2 readily infected well-differentiated proximal airway cells leading to viral replication and gene expression as indicated by induction of SARS-CoV-2 Nucleoprotein gene (N gene) RNA in infected samples. N gene abundance, reflective of either viral mRNA expression or a result of genome replication increased 750-fold at 2dpi and declined at 3dpi, however, the later decline was not statistically significant. Viral N gene expression was undetectable in corresponding mock cultures at all 3 time points (Figure 1B).

**Fig. 1.**
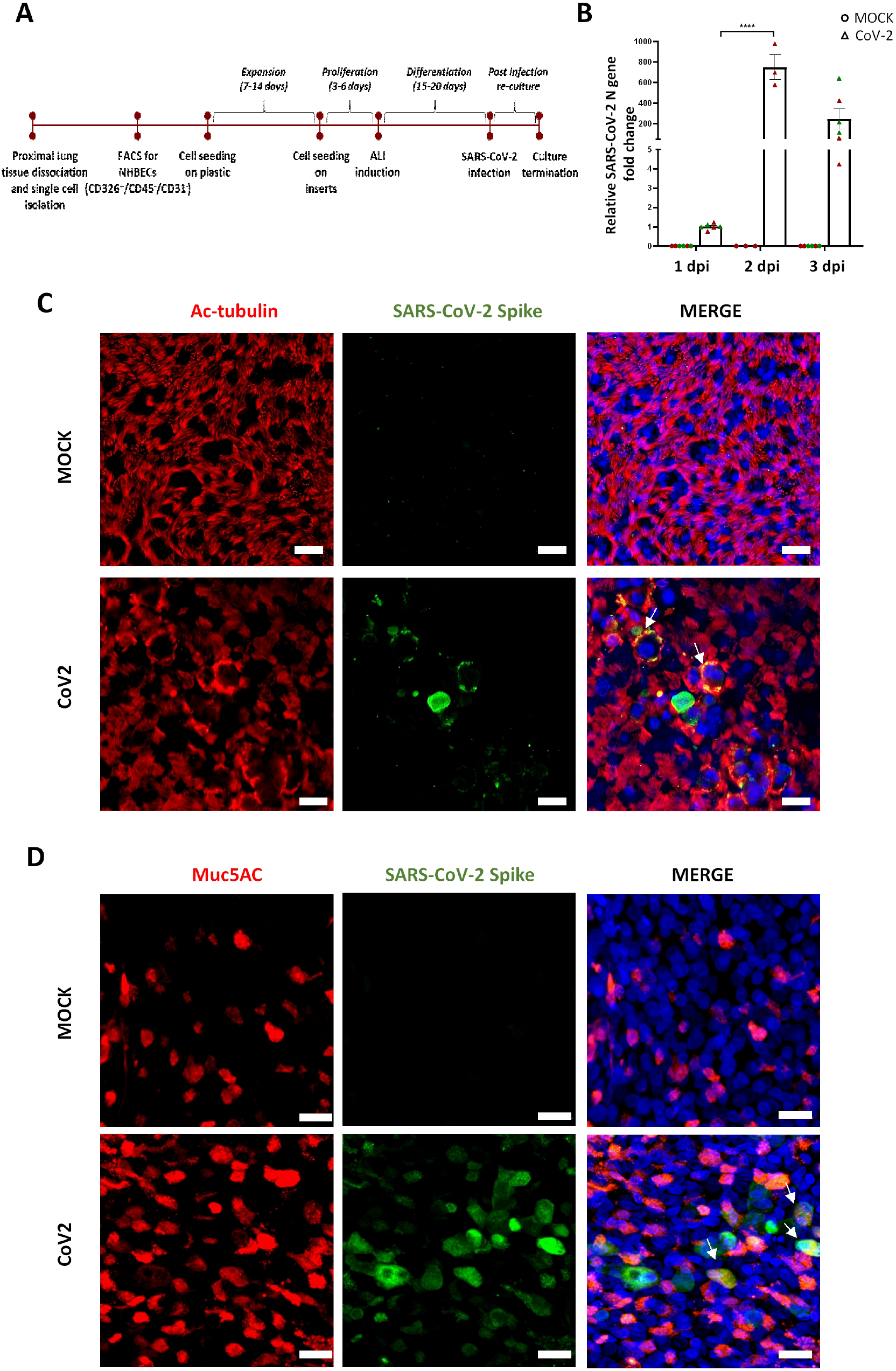
SARS-CoV-2 infects and replicates within normal human proximal airway cells. **(A**) Workflow for establishment of human proximal airway ALI cultures and their infection with SARS-CoV-2. **(B)** ALI cultures of proximal airway epithelial cells are susceptible to SARS-CoV-2 infection which peaked at 2dpi; n= 3 to 6 cultures from 2 independent donors. Circles indicate mock cultures and triangles indicate SARS-CoV-2 infected cultures. Red and green colors indicate cultures from separate donors. Data were analyzed using Two-Way ANOVA with Sidak’s post-hoc correction and represented as fold change for individual cultures ± SEM relative to mean N gene expression at 1dpi ****p<0.0001. 2 days after SARS-CoV-2 infection, viral spike protein (green) was found to heterogeneously co-localize with **(C)** acetylated tubulin positive ciliated cells (red) and **(D**) a proportion **of** Muc5AC positive goblet cells (red). Scale bar = 20μm. Arrows indicate colocalization of markers.

Further analysis of infected cultures revealed that SARS-CoV-2 infection of proximal airway ALI cultures was heterogenous. SARS-CoV-2 predominantly targeted ciliated cells, evidenced by colocalization of SARS-CoV-2 viral capsid “spike” protein with the ciliated cell marker, acetylated tubulin (Figure 1C). This observation is congruent with previous reports of infection of ciliated cells by SARS-CoV (*11*) and recent reports of infection of ciliated cells by SARS-CoV-2, in *in vitro* cultures of proximal airways (*12, 13*) as well as COVID-19 patient biopsies (*12*). A number of studies have described the expression of the SARS-CoV-2 entry receptor, angiotensin-converting enzyme2 (ACE2) in the respiratory tracts (*12, 14–16*). In the nasal passages and large airways highest levels of ACE2 expression have been reported in ciliated cells (*12, 15, 17*). However, we also observed SARS-CoV-2 infection within a small proportion of goblet cells, expressing the mucin, Muc5ac (Figure 1D).

### SARS-CoV-2 infects and replicates within human alveolar type 2 cells

A number of recent studies have utilized various cell lines, primary epithelial cultures of nasal and proximal airway epithelium as well as organ on chip methods to study SARS-CoV-2 infection of the upper airways, the initial site of entry and infection (*12, 18–20*). However, there is a lack of a tractable system to model SARS-CoV-2 infection in the distal gas-exchange region of the lung, the alveolus, which represents the site of severe pathology and pneumonia. Recent single cell transcriptome studies have shown that ACE2, the host cell surface receptor for SARS-CoV-2 attachment and infection, is predominantly expressed by alveolar type 2 (AT2) cells (*12, 16*). AT2 cells are bifunctional cells that serve both as progenitors that contribute to epithelial maintenance in addition to fulfilling specialized functions such as surfactant production (*5, 21*). We established a 3D organoid culture model of the human alveoli to model SARS-CoV-2 infection of the distal lung in *in vitro* (Figure 2A). This is the first study to report the development of a primary *in vitro* model of the adult human alveoli to study the pathogenesis of SARS-CoV-2 infection. Using this system, primary epithelial cells isolated from normal human distal lung tissue were cultured in the presence of stromal support cells leading to the generation of alveolar organoids (*5*). Organoid cultures were exposed to SARS-CoV-2 (1×10^4 TCID50 per well) and harvested for analysis 1-3 dpi. We found that intact organoids were refractory to viral infection but gentle physical and enzymatic disruption was permissive for viral infection and replication, as assessed by relative N gene expression (Figure 2B). Disruption of organoids exposed the apical cell surfaces of cells that otherwise face inwards towards the air-filled lumen in the 3D organoid structure, thus presumably enabling the virus to access its entry receptor, ACE2. Study of viral infection kinetics from 1dpi to 3dpi demonstrated that viral infection peaked at 2dpi. N gene expression at 2dpi was 7-fold higher compared to the mean infection at 1dpi and declined significantly by 3dpi. Viral N gene expression was below the detection limits in corresponding mock cultures at all 3 time points (Figure 2C).

**Fig. 2.**
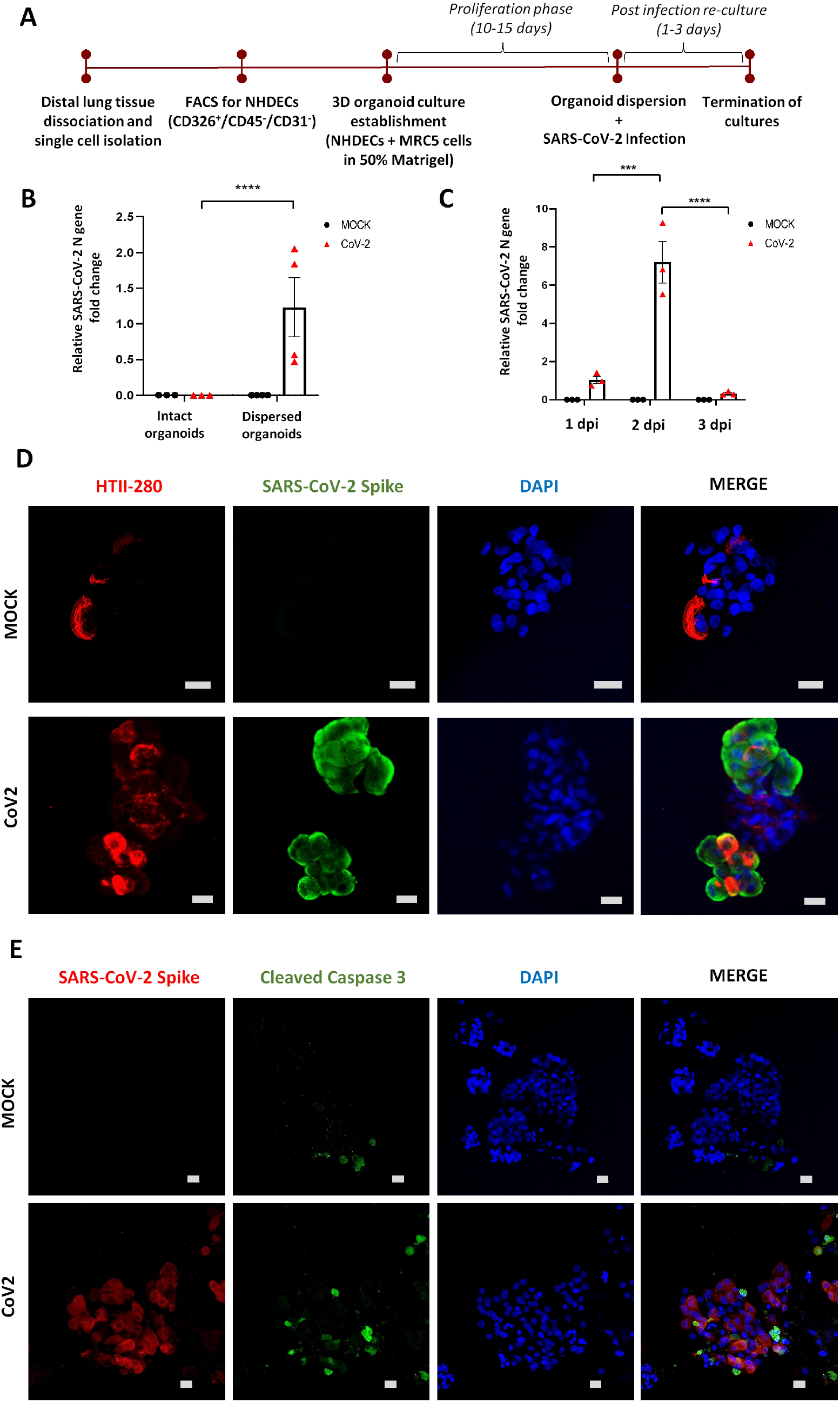
SARS-CoV-2 infects and replicates within normal human distal alveolar organoids. **(A)** Workflow for establishment of human distal alveolar organoid cultures and their infection with SARS-CoV-2. **(B)** Enzymatic and mechanical disruption of alveolar organoids to expose the apical surface was essential for robust infection, as evaluated by detection of SARS-CoV-2 N gene abundance; n = 3 to 4 organoid cultures established from 2 different donor samples for each condition. **(C)** Viral infection levels peaked at 2 days post infection; n=3 organoid cultures. Data were analyzed using Two-Way ANOVA with Sidak’s post-hoc correction and represented as fold change for individual cultures ± SEM relative to mean N gene expression at 1dpi. ****p=0.0002 for **(B)** and ***p=0.0009 ****p<0.0001 for **(C)**. **(D)** Viral infection was assessed at 2dpi by antibody against SARS-CoV-2 “spike” protein (green) and colocalized with AT2 cell marker HTII-280 (red) **(E)** Infected alveolar organoids (red) also demonstrated signs of cellular apoptosis at 3dpi, indicated by positive staining for cleaved caspase-3 (green); scale bar= 20μm.

We used immunofluorescence confocal microscopy to confirm infection of organoid cultures with SARS-CoV-2. Mock and infected alveolar organoids were stained for the well-known AT2 cell marker, HTII-280 and SARS-CoV-2 spike protein. Organoid cultures showed abundant expression of HTII-280 and in infected cultures, viral spike protein localized to the HTII-280^+^ AT2 cells (Figure 2D). By 3dpi, infected alveolar organoids exhibited significantly enhanced staining for the apoptotic marker, cleaved caspase-3 compared to mock cultures. Interestingly only a proportion of the apoptotic cells were infected, thus demonstrating that SARS-CoV-2 induced a cytopathic effect on infected AT2 cells as well as neighboring uninfected cells (Figure 2E, Supplementary Figure 1). SARS-CoV-2 induced apoptosis of neighboring uninfected epithelial cells is suggestive of the potential for a non-cell autonomous effect on alveolar epithelial integrity.

Taken together, these data demonstrate that AT2 cells are susceptible to SARS-CoV-2 infection and suggest that infection triggers both cell-autonomous and non-cell-autonomous apoptosis that may contribute to alveolar injury. We conclude that 3D alveolar organoid cultures serve as a robust platform for studying the effect of SARS-CoV-2 infection on adult distal lung alveolar epithelium.

### 3D alveolar organoid cultures as a tool to study host response to SARS-CoV-2 infection

Having established a 3D alveolar organoid model that enables robust infection of AT2 cells by SARS-CoV-2, we further evaluated the utility of this system to study host pathogen responses. Distal organoid cultures were infected with SARS-CoV-2 and cells were harvested 2 dpi for transcriptomic analysis via global RNA-Sequencing. Transcript profiles were compared between SARS-CoV-2 infected and mock organoid cultures. Heatmap (Figure 3A) and volcano plots (Figure 3B) of differentially expressed genes revealed detection of high levels of SARS-CoV-2 viral RNA such as coding sequences for *Virus_N, virus_ORF1ab, virus_ORF3a*b, further confirming SARS-CoV-2 genome replication and/or gene transcription in organoid cultures. Furthermore, infected organoids showed robust induction of host response genes including cytokines such as *IFNB1* and antiviral response genes *OAS1, OAS2, ISG15* and *MX1*. We did not see a significant change in expression of the AT2 cell marker, *SFTPC*, or host genes involved in viral infection such as *ACE2* and *TMPRSS2*. Analysis of differentially regulated canonical pathways revealed that Interferon (IFN) signaling was the most upregulated pathway. Other upregulated pathways included NF-kB activation, TLR signaling and IL1 signaling (Figure 3C). On the other hand, antigen presentation, Th1 and Th2 activation pathways were downregulated (Figure 3D). Taken together, these data indicate that SARS-CoV-2 infection induces significant alteration of innate immune response genes in alveolar cells without the participation of recruited immune cells. We speculate that the epithelial innate immune response may provide activating signals leading to global activation of the host immune response and provide therapeutic targets for mitigation of uncontrolled lung inflammation and adverse patient outcomes.

**Fig. 3.**
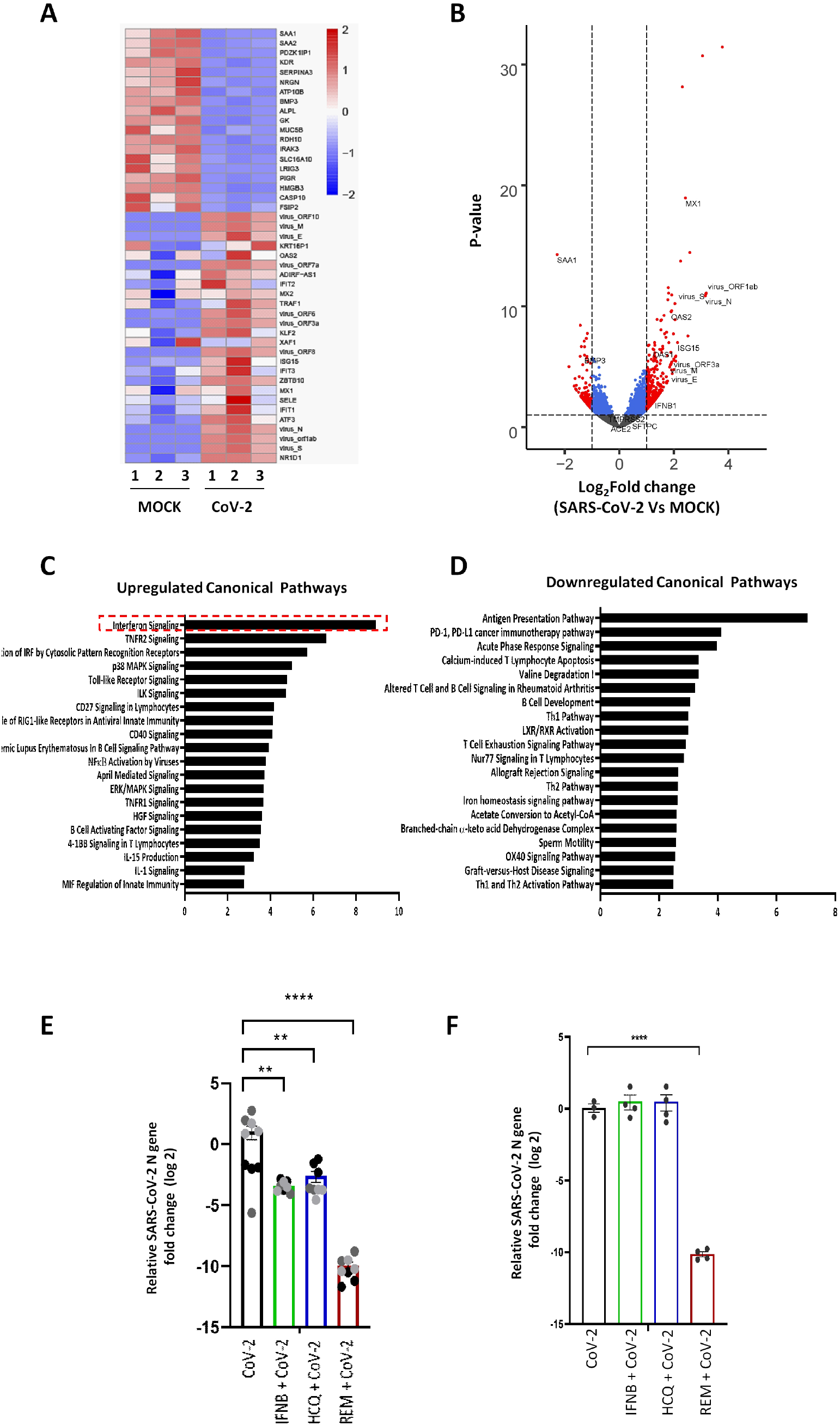
Primary human lung epithelial models for study of SARS-CoV-2 induced host response and drug validation. **(A)** Heatmap showing RNA seq analysis of differentially expressed genes at 2 dpi. (**B)** Volcano plot of gene expression changes in SARS-CoV-2 infected vs mock cultures defined by p-value and >2-fold change. Several viral genes such as *Virus_N, virus_ORF1ab, virus_ ORF3a* were detected in the infected samples. Cytokines such as *IFNB1*, and antiviral response genes *OAS1, OAS2, ISG15 and MX1* were significantly upregulated. **(C)** The most upregulated canonical transcriptional pathway in SARS-CoV-2 infected alveolar cultures was IFN signaling pathway. In addition to TLR and NFKB signaling. **(D)** Downregulated canonical transcriptional pathways include antigen presentation and Th1 and Th2 activation pathways. **(E)** Pre-treatment of alveolar organoid cultures with IFNB1, Hydroxychloroquine and Remdesivir significantly reduced viral replication. The effect of Remdesivir on viral replication was more pronounced than that of IFNB1 or hydroxychloroquine. Data are represented as log2fold change for individual cultures ± SEM, normalized to mean infection and analyzed using One-Way ANOVA with Tukey’s post-hoc test **p=0.0012 for IFNB1, **p=0.0044 for HCQ and ****p<0.0001 for Remdesivir. The 3 different colors indicate cultures from different biological replicates. **(F)** Pre-treatment of proximal ALI cultures with Remdesivir significantly reduced viral infection/replication. Pre-treatment with Hydroxychloroquine and IFNB did not have an effect on viral replication; n= 3-4 independent cultures. Data are represented as log_2_fold change for individual cultures, normalized to mean infection and analyzed using One-Way ANOVA plus Tukey’s posthoc test ****p<0.0001.

### Organoid and ALI culture systems as a tool to screen therapeutic targets against SARS-CoV-2 infection

Given that animal models such as mice are not natural hosts to SARS-CoV-2 infection, there is a vital need for development of alternate pre-clinical models that closely recapitulate the human lung for screening potential therapeutic agents that target SARS-CoV-2 infection and replication. We utilized the 3D alveolar organoid and proximal airway ALI culture systems to study the effect of a selected panel of drugs which included the known anti-viral cytokine, IFNB1 and investigational drugs for COVID-19 treatment, Remdesivir (*22, 23*) and Hydroxychloroquine (*24, 25*). Treatment of 3D alveolar organoid cultures with IFNB1 lead to a 3.2-log reduction in viral N gene RNA compared to control infected organoids. Hydroxychloroquine led to an overall 2.4-log reduction in viral N gene expression compared to average infection in untreated organoids (Figure 3E). However, variable effects of hydroxychloroquine were observed on viral replication/gene expression that were donor epithelium-dependent (Supplementary Figure 2). Remdesivir showed the strongest effect on viral replication in alveolar organoids, resulting in a 9-log decrease in viral N gene expression compared to the average infection in untreated organoids (Figure 3E). This effect was consistent irrespective of donor origin of epithelial cells confirming it as a direct acting antiviral (DAA) agent targeting viral specific RNA polymerase (Supplementary Figure 2). IFNB1 and Hydroxychloroquine did not show a significant effect on viral N gene RNA in proximal ALI cultures. Proximal airway ALI cultures used for this study were derived from separate donors compared to alveolar organoid cultures. The lack of response to IFNB1 may reflect differences in disease susceptibility observed in different COVID-19 patients. However, further analysis is warranted to confirm this observation. However, Remdesivir resulted in a 10.2-log reduction in viral N gene RNA abundance, confirming its effect as a DAA (Figure 3F). Taken together, our data show that 3D alveolar organoid models and proximal ALI cultures represent a highly relevant preclinical tool to assess SARS-CoV-2 infection and replication, and serve as a sensitive platform for drug screening and validation.

Overall, our study provides a novel model to investigate SARS-CoV-2 infection along the proximal-distal axis of the human lung epithelium. SARS-CoV-2 infection is accompanied by a cell-autonomous proinflammatory response and viral infection and/or replication is strongly suppressed by the candidate drug, Remdesivir. This platform can help better understand the pathogenesis of COVID-19. Furthermore, our platform provides a relevant preclinical model for rapid screening of drugs against COVID-19 and future emergent respiratory pathogens.

## Acknowledgments

We would like to thank the Cedars-sinai Applied Genomics, Computation and Translational Core for their help with performing bulk RNAseq. The following reagents were obtained through BEI Resources, NIAID, NIH: Monoclonal Anti-SARS-CoV S Protein (Similar to 240C), NR-616; SARS-Related Coronavirus 2, Isolate USA-WA1/2020, NR-52281.

## Funding

This work was supported by National Institutes of Health (5RO1HL135163-04, PO1HL108793-08, T32HL134637), Celgene IDEAL Consortium and Parker B. Francis Fellowship Program.

## Author contributions

A.M. and B.K. performed experiments. A.M. and B.R.S. designed studies and wrote the manuscript. G.G.Jr., S.B., C.S., A. P., P.P., J.S.D.A., and C.G.D.A, prepared samples. C.Y. analyzed RNA seq data. A.M., C.K., B.G., B.R.S. and V.A. interpreted results. J.K.K. and D.A.P. provided IFNB1 and guidance for drug validation experiments. B.R.S. and V.A. supervised.

## Competing interests

Authors declare no competing interests.

## Notes

### Competing Interest Statement

The authors have declared no competing interest.

