## Supplementary Figures for "SARS-CoV-2 infection of primary human lung epithelium for COVID-19 modeling and drug discovery"

**Abstract:** Coronavirus disease 2019 (COVID-19) is the latest respiratory pandemic resulting from zoonotic transmission of severe acute respiratory syndrome-related coronavirus 2 (SARS-CoV-2). Severe symptoms include viral pneumonia secondary to infection and inflammation of the lower respiratory tract, in some cases causing death. We developed primary human lung epithelial infection models to understand responses of proximal and distal lung epithelium to SARS-CoV-2 infection. Differentiated air-liquid interface cultures of proximal airway epithelium and 3D organoid cultures of alveolar epithelium were readily infected by SARS-CoV-2 leading to an epithelial cell-autonomous proinflammatory response. We validated the efficacy of selected candidate COVID-19 drugs confirming that Remdesivir strongly suppressed viral infection/replication. We provide a relevant platform for studying COVID-19 pathobiology and for rapid drug screening against SARS-CoV-2 and future emergent respiratory pathogens.

**One Sentence Summary:** A novel infection model of the adult human lung epithelium serves as a platform for COVID-19 studies and drug discovery.

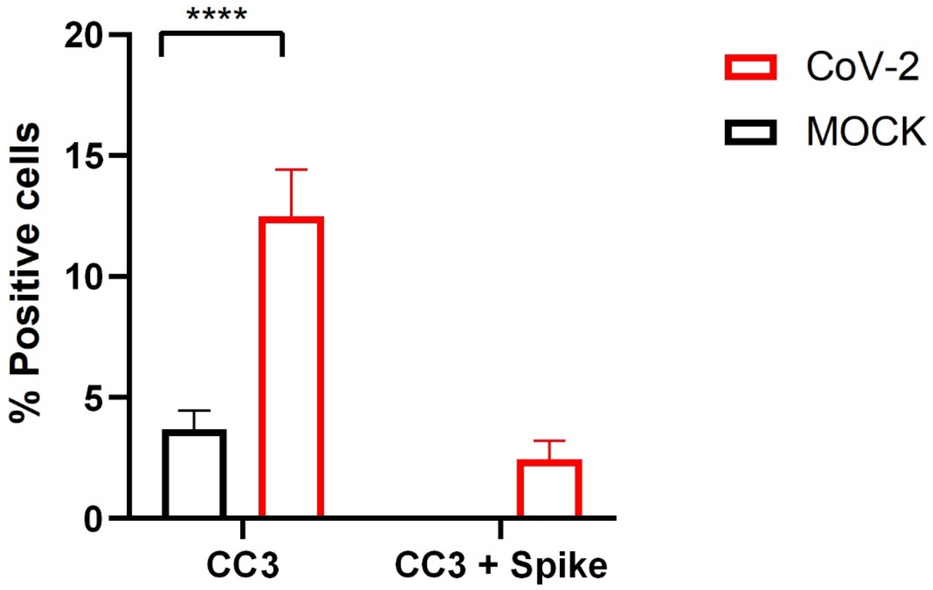


**Fig. S1. SARS-CoV-2 induces cell-autonomous and non-cell-autonomous apoptosis of AT2 cells.**

Number of cleaved-caspase 3 positive cells was significantly higher in SARS-CoV-2 infected alveolar organoids compared to mock alveolar organoids. Only a percentage of the apoptotic cells were positive for viral spike protein indicating that SARS-Co-V2 infections induces apoptosis of neighboring cells in addition to infected cells. Spike protein staining was indetectable in mock cultures. N= 7 different images of alveolar organoids for each condition. Data are represented as mean percentage of cells staining positive for target protein ± SEM and analyzed using Two-Way ANOVA plus Sidak's post-hoc test ****$p<0.0001$.

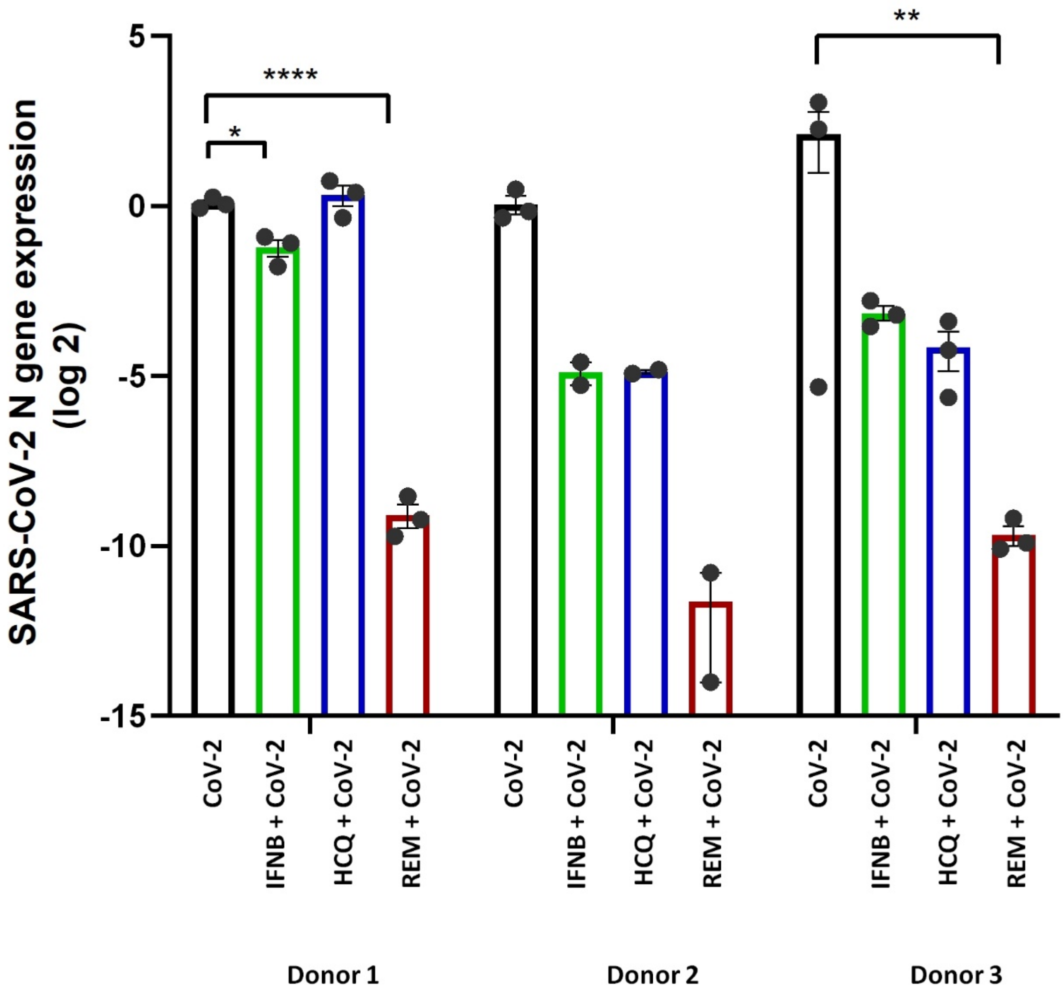


**Fig. S2. Effect of drugs on 3D alveolar organoid cultures from individual biological replicates.**

Effect of treatment of IFNB1, Hydroxychloroquine and Remdesivir on 3D organoid cultures from 3 different donors showed variability for IFNB1 and hydroxychloroquine effect amongst donors; Variability among donors for hydroxychloroquine effect was higher. However, Remdesivir showed consistently strong inhibition of viral replication/infection irrespective of donor origin of cells. N= 2 to 3 cultures from 3 independent biological replicates. Data are represented as $log_2$fold change for individual cultures ± SEM, normalized to mean infection for individual biological replicates and analyzed using One-Way ANOVA plus Tukey’s post-hoc test within individual biological replicate Donor 1: *$p=0.021$ ****$p<0.0001$; Donor 3: ** $p=0.0024$ ****$p<0.0001$.
